# A Sustainable Approach to Natural Food Coloring: Alginate-Based Encapsulation and Stabilization of Beetroot Betalain

**DOI:** 10.64898/2026.01.15.699826

**Authors:** Amruth N Murthy, Kavana N Murthy, M L Harshith, Y Channarushabendra, B Trilok Chandran

## Abstract

Betalains are natural, water-soluble pigments that are chiefly extracted from Beta vulgaris(beetroot) and have been widely researched as natural food colorants attributed to their natural coloring ability and antioxidant properties. On the other hand, their use is restricted by their poor stability response against exposure to natural elements like light, temperature fluctuations, oxygen, and changes in pH, which causes rapid decomposition of pigments and loss of coloration. The scope of this research is to improve the stability response of beetroot-extracted betalain against natural element exposure by employing alginate encapsulation and co-encapsulation techniques with food-grade components. Betalain extracts for this research are conducted through a method of methanolic extraction, and the concentration of the pigment is measured by UV-Visible Spectrophotometry at a wavelength of 535 nm. The encapsulation method is performed through the ionotropic gelation method of sodium alginate and calcium chloride, while the co-encapsulation method is performed by embedding L-ascorbic acid, which is naturally recognized as a non-toxic natural antioxidant agent, within the sodium alginate matrix. The above systems were assessed for color retention upon storage, pH stability at an acidic pH of 3 and basic pH of 9, and controlled release properties in an aqueous environment. The results showed that alginate encapsulation of the betalains increased the retention of color stability and pH stability significantly when compared to the free pigment. Alginate co-encapsulation of the betalains performed better in color stability and controlled release properties because of the reduced diffusion caused by the addition of ascorbic acid. Generally, alginate co-encapsulation of the betalains offers a sustainable and food-friendly approach for the stabilization of the betalains for application in food colorants and controlled release systems.

## I. Introduction

A major reason that has escalated the demand for natural food colorants has been the increasing preference for clean label and natural-origin food ingredients among consumers in recent years [1], [19]. Although natural food colorants are expensive and less stable compared to artificial ones, their preference has escalated because artificial colored foods have been perceived as hazardous to consumers’ health and are subjected to strict regulations in many countries. Plant-based colors can be considered a preferable alternative because of their biodegradability and biocompatibility in addition to their functional properties. Betalains are concerned with pigments containing nitrogen and are water-soluble; they are responsible for the red-violet to yellow coloration of plants of the order Caryophyllales. Betalains are abundantly found in beetroot (Beta vulgaris), which is one of the natural sources of red-colored compounds like betalains exhibiting considerable properties as antioxidants, anti-inflammatory compounds, and chemoprotective agents [1], [18], [19]. Due to this property, there is a growing demand for the use of betalains in food coloration, nutraceuticals, and other pharmaceuticals. Despite their promising qualities, betalains’ poor stability under processing and environmental conditions severely restricts their widespread industrial use. Betalain pigments are extremely sensitive to light, high temperatures, oxygen, and pH changes, especially in alkaline environments. During food processing and storage, these factors hasten pigment degradation, resulting in color fading and a loss of bioactivity, which lowers the food’s commercial viability [2], [5]. Sensitive bioactive compounds can now be effectively shielded from unfavorable environmental conditions by encapsulation. Encapsulation can increase stability, extend shelf life, and allow for controlled release of the active ingredient by enclosing pigments in a protective polymeric matrix. Due to its natural origin, non-toxicity, biodegradability, and approval for use in food applications, sodium alginate is a popular encapsulating material. Alginate can be used to encapsulate water-soluble pigments like betalains because it ionically cross-links with calcium ions to form stable gel beads [8], [15]. Co-encapsulation adds additional protection to pigment protection through the inclusion of stabilizing compounds in the encapsulation matrix. Ascorbic acid, a biological antioxidant, has been proposed to non-enzymatically reduce betalain degradation by scavenging reactive oxygen molecules [7], [10]. It is therefore expected that the combination of alginate encapsulation and ascorbic acid co-encapsulation would result in a synergistic stabilization effect. In this investigation, the aim is to isolate and determine the concentration of betalain from beetroot and to explore the efficacy of alginate encapsulation and ascorbic acid coencapsulation as means to enhance betalain stability. The significance and novelty of this investigation are the assessment and performance analysis of the stability and controlled release properties of the encapsulated as well as the co-encapsulated betalain through the use of the sustainable and biodegradable process, its use as natural food coloring.

## II. Problem Statement

The growing requirement for natural and clean-label food items has thus increased the requirement for finding natural substitutes for synthetic food colorants. Beetroot (Beta vulgaris) is one source of betalain pigments possessing impressive red color along with other beneficial health properties, which make them useful for natural food coloration purposes. The use of beetroot betalain is greatly hindered by its natural instability when subjected to various process treatments. Betalains are very sensitive to certain conditions, including light, temperature, the presence and level of oxygen, and pH, especially an alkaline environment. This results in the rapid degradation and loss of the bioactivities exhibited by the pigments when exposed to the conditions cited above, thereby being responsible for the lackluster stability and color properties displayed by conventionally extracted betalains. Although many stabilization strategies have been investigated, many of the current strategies use artificial additives or intricate processing techniques that are not entirely compliant with sustainability and clean-label regulations. A straightforward, environmentally friendly, and food-grade stabilization method that can successfully safeguard betalain pigments without sacrificing safety or legal compliance is obviously needed [6], [15]. Additionally, only a small number of studies have thoroughly assessed the use of natural polymer-based encapsulation in conjunction with antioxidant co-encapsulation to enhance betalain stability, specifically in terms of color retention, pH stability, and controlled release behavior. By creating an alginate-based encapsulation and co-encapsulation system with ascorbic acid as a natural stabilizer, this study fills this research gap by improving the industrial applicability of beetroot betalain as a natural food colorant in a sustainable and food-compatible manner.

## III. Materials and Methods

### A. Materials

Raw beetroot specimens containing beta vulgaris pigment were obtained from a local market. Sodium alginate was used as an encapsulant, and calcium chloride (CaCl2) was used as the cross-linker. Methanol, an analytical grade solvent, was used as the extractant, and ascorbic acid functioned as the natural antioxidant in the process of co-encapsulation. Distilled water was used in all experiments. Hydrochloric acid (0.1 N) and sodium hydroxide (0.1 N) solutions were used as pH adjustment agents. The apparatus and equipment utilized in this experiment included a blender for homogenization, centrifuge, Whatman No.1 filter paper, UV-Visible spectrophotometer, quartz cuvettes with a path length of 1 cm, magnetic stirrer, syringe for preparing the beads, pH indicator paper, refrigerator at 4 °C, and laboratory glassware.

**Fig. 1.**
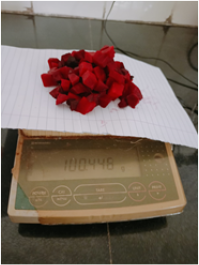
Washed and Peeled Beetroot

### B. Extraction and Quantification of Betalain

Betalain pigment was extracted from fresh beetroot using a methanolic extraction method. The beetroot samples were carefully washed, peeled, and cut into small pieces. A specific amount of beetroot was blended with methanol to create a uniform slurry. The mixture was filtered through Whatman No.1 filter paper to remove solid particles. It was then centrifuged at 5000 rpm for 10 minutes to obtain a clear liquid. The betalain extract was collected and stored in amber containers at 4 °C to avoid light-induced degradation. To quantify betalain, we used a UV-Visible spectrophotometer. We measured the absorbance of the appropriately diluted samples at 535 nm [3], [5], [7], which is the maximum absorption wavelength of betacyanin. Methanol served as the blank, and we calculated the betalain concentration using Beer-Lambert’s law.

**Fig. 2.**
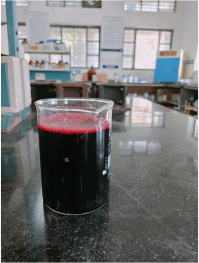
Crude betalain extract

### C. Preparation of Alginate Beads

#### 1) Encapsulation of Betalain(Group A)

A 2% (w/v) solution of sodium alginate was prepared by dissolving the appropriate amount of sodium alginate in distilled water, stirring continuously. Equal parts of betalain extracts and solution of sodium alginate were mixed to form a uniform solution. The solution was transferred to a syringe and dropped slowly into a solution of 0.1 M CaCl2 from a fixed position, which resulted in the process of ionic gelation [6], [8]. The Ca alginate beads formed immediately. The beads were allowed to settle in the CaCl2 solution for 15 minutes and were then removed, rinsed in distilled water to remove any excess Ca2+ ions, and referred to as betalain beads (Group A).

**Fig. 3.**
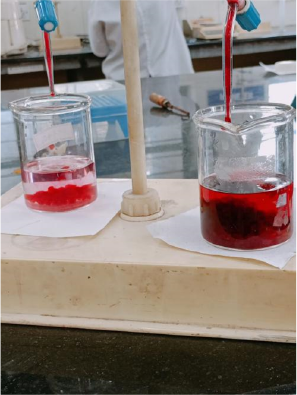
Encapsulated and Co-encapsulated Betalain Beads

#### 2) Co-Encapsulation of Betalain with Ascorbic Acid (Group B)

For co-encapsulation, a solution of ascorbic acid was added to the betalain extract solution prior to mixing with alginate solution. The mixture of betalain and ascorbic acid solution was then further mixed together with a solution of sodium alginate. The combined solution was then dropped slowly into a solution of calcium chloride using ionic gelation. The formed beads were then cross-linked for 15 minutes and later labeled as co-encapsulated betalain beads (Group B).

### D. Color Retention Study

The storage stability of free, encapsulated, and coencapsulated betalain was investigated. The samples were wrapped with aluminum foil and stored at 4 °C. In the cases of encapsulated/co-encapsulated samples, equal numbers of beads were considered for each experiment. Color intensity was determined at 0, 24, and 48 hours, by disrupting the beads in distilled water to release the pigment. The absorbance was then measured at 535 nm using a UV–Visible spectrophotometer. Color retention percentage was determined by comparing each time absorbance value with the initial absorbance value.

### E. pH stability Study

In both acidic and alkaline environments, betalain’s pH stability was evaluated. 0.1 N hydrochloric acid and 0.1 N sodium hydroxide were used to create buffer solutions with pH values of 3 and 9, respectively. While free betalain extract was examined independently, encapsulated and co-encapsulated beads were submerged in the corresponding buffer solutions. Beads were crushed in the buffer medium at predefined intervals while the samples were incubated at room temperature.

**Fig. 4.**
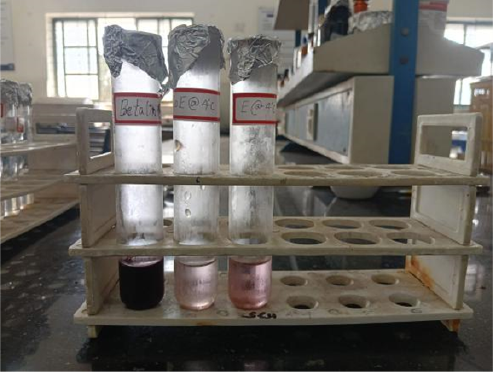
Color Retention Study

The percentage of absorbance that was retained in relation to the initial value was used to express pH stability. Absorbance was measured at 535 nm.

**Fig. 5.**
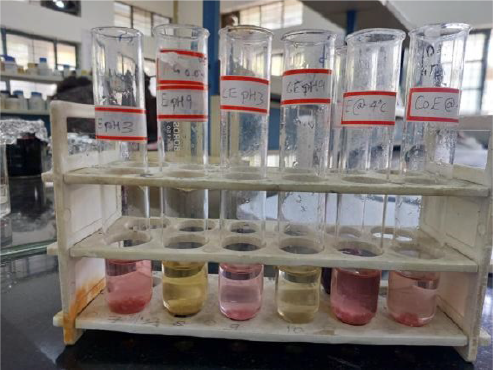
pH Stability Study

### F. Controlled Released Study

The controlled release study of the betalains was performed using the beads made from the alkaline soluble component of the alginate, using distilled water as the dissolution media. Equal numbers of beads with and without the encapsulated sample were used and placed in separate tubes containing a fixed amount of distilled water at room temperature. At set intervals, an amount of the dissolution media was removed and was followed by the immediate addition of an equivalent amount of distilled water. Spectrophotometric measurement was performed at 535 nm, and the cumulative amount of dissolution was determined relative to the net absorbance at set intervals, corresponding to the total potential amount at complete dissolution.

**Fig. 6.**
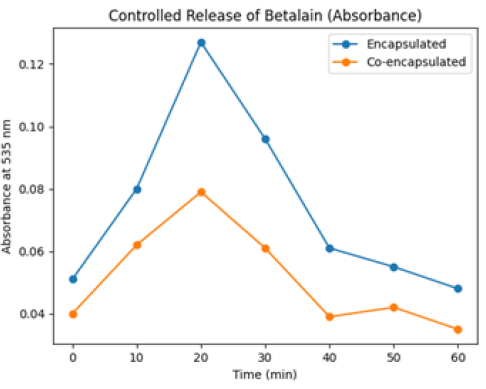
Controlled Released Study

## IV. Results and Discussion

### A. Colour Retention Analysis

The color retention behavior of free, encapsulated, and co-encapsulated beetroot betalain was studied by measuring absorbance at 535 nm as a function of storage time at 4 °C. Absorbance decreased noticeably over time for all samples, indicating slow but progressive degradation of the pigments. However, the kinetics and overall color retention differed significantly among the three systems. Free betalains underwent the most pronounced and rapid losses of absorbance, reflecting poor stability and high susceptibility to deteriorative changes in the environment during storage. By contrast, encapsulated betalain exhibited better color retention, confirming the protective action of the alginate matrix. By acting as a physical barrier, the calcium alginate beads reduced the amount of light, oxygen, and other destabilizing elements that could reach the pigment. This encapsulation improved pigment stability by slowing the rate of oxidative and photoinduced degradation [7], [10]. Co-encapsulated betalain beads showed the best color retention, maintaining a significantly higher percentage of their initial absorbance during the storage period. Alginate encapsulation and ascorbic acid’s antioxidant properties work together to improve stability [6], [15], [16]. Ascorbic acid minimizes pigment degradation by efficiently scavenging reactive oxygen species and lowering oxidative stress within the bead matrix. The general pattern was coencapsulated betalain ¿ encapsulated betalain ¿ free betalain, which amply illustrates the co-encapsulation method’s superiority for maintaining betalain color intensity.

**Fig. 7.**
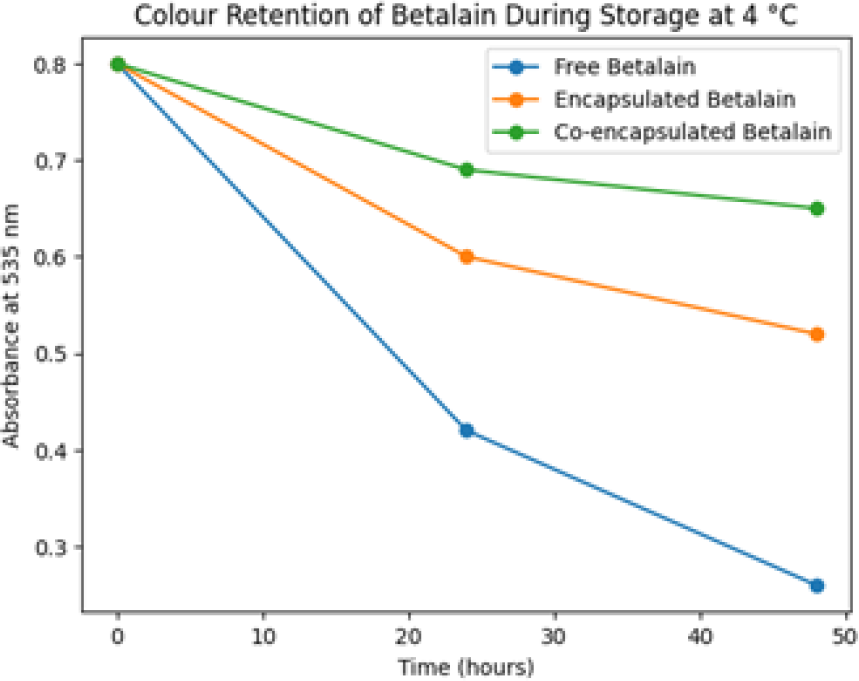
Color Retention Profile

### B. pH Stability Behavior

The pH stability of the encapsulated and co-encapsulated betalain solutions was tested using the acidic pH 3 and basic pH 9 solutions, and the stability of the betalains was measured in terms of the absorption of light by the solutions at 535 nm. The data clearly shows the effect of pH on the stability of the betalains [2], [18], [19], with both the encapsulated and coencapsulated betalains having a relatively high absorption in the case of the acidic solution compared to the basic solution, and this aligns with the chemical stability of the betalains in the acidic solution. Betalain co-encapsulated was found to have higher stability at pH 3, especially when the incubation time was prolonged, indicating a synergistic protection against degradation from ascorbic acid. The alginate matrix itself also played a part in enhancing stability by effectively suppressing diffusion and hence minimizing interactions with the external environment. On the contrary, the alginate matrix was found to cause a great reduction in the absorption values in both systems at higher pH, clearly illustrating the high sensitivity of betalain to alkaline degradation [2], [5]. Despite the protection offered by encapsulation, the prevention of degradation by ascorbic acid was not as effective. Diffusion-controlled release of betalain from the alginate beads, rather than pigment regeneration, can be the cause of an increase in absorbance seen at later time points for some samples. The dynamic balance between pigment degradation and slow diffusion from the encapsulated system, which is frequently seen in polymer-based delivery matrices, is reflected in this release-related absorbance variation.

**Fig. 8.**
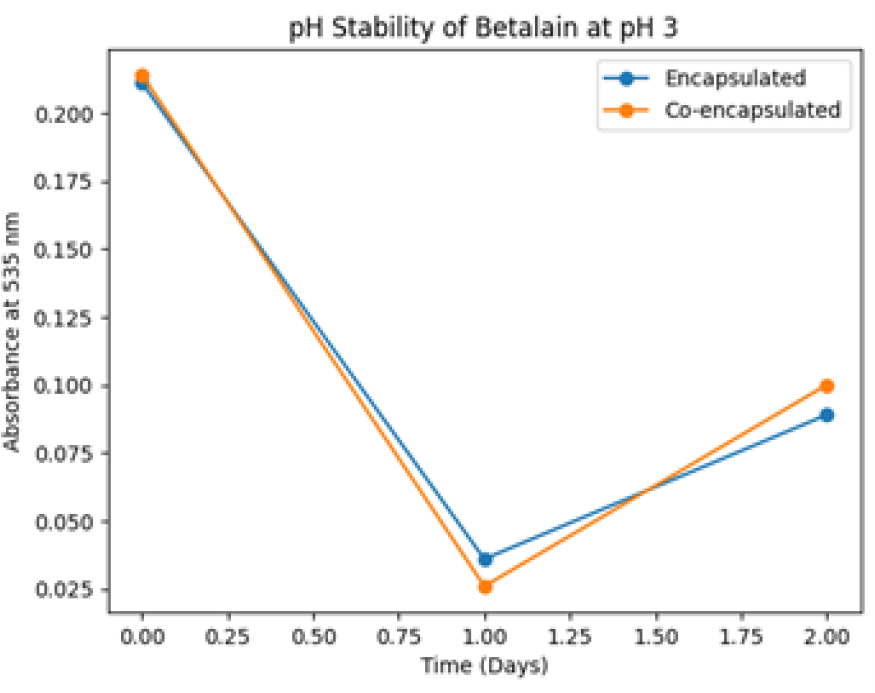
Beads Stability at pH3

### C. Controlled Release Profile

The controlled release of betalain from encapsulated and co-encapsulated alginate beads was measured in aqueous conditions by tracking the absorbance of 535 nm for 60 minutes. The two systems portrayed a positive increase in absorbance, representing the rapid release of the surface-bound pigment, often referred to as the initial burst release. The rate of release of the encapsulated betalain was faster, where the absorbance reached its peak within a shorter time, hence having low resistance during diffusion into the alginate beads [15], [16]. By contrast, the co-encapsulated beads showed a relatively slower and more sustained rate of pigment release. The presence of ascorbic acid is possibly accountable for compaction of the matrix and contraction of its pore size, which in turn caused greater diffusion resistance and delayed rate of pigment release. After the completion of rapid diffusion, a rather gradual loss of absorbance was noticed in both the formulations. The sustained release properties exhibited by beaded preparations of co-encapsulated materials are especially beneficial in food applications where the stability of color and controlled distribution of pigments are of prime importance. The achievements of this research show that coencapsulation based on alginate is very promising and offers not only stability of betalain, but it is also useful in controlled release technology.

**Fig. 9.**
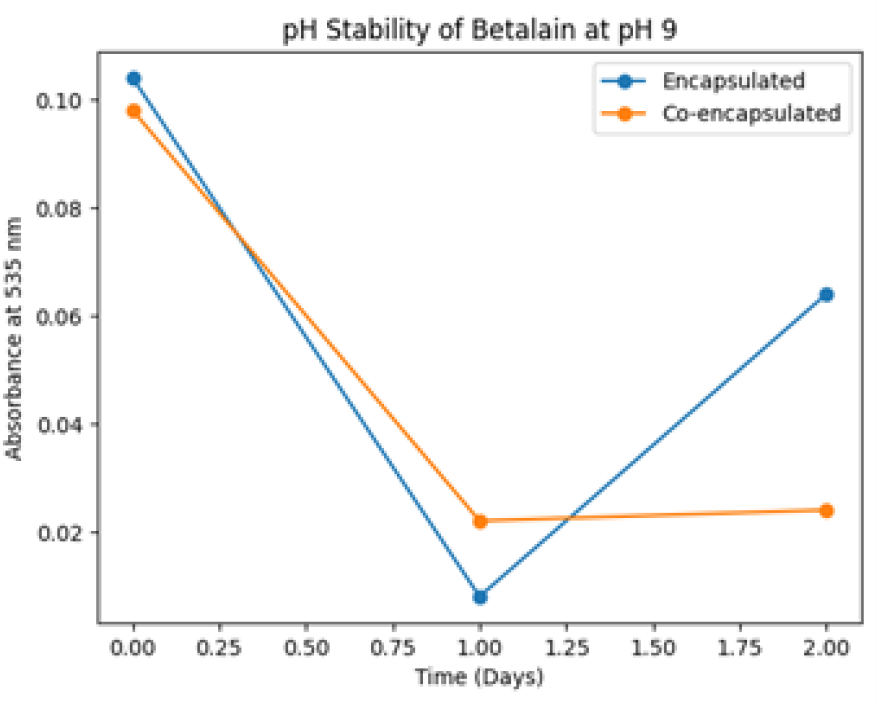
Beads stability at pH9

**Fig. 10.**
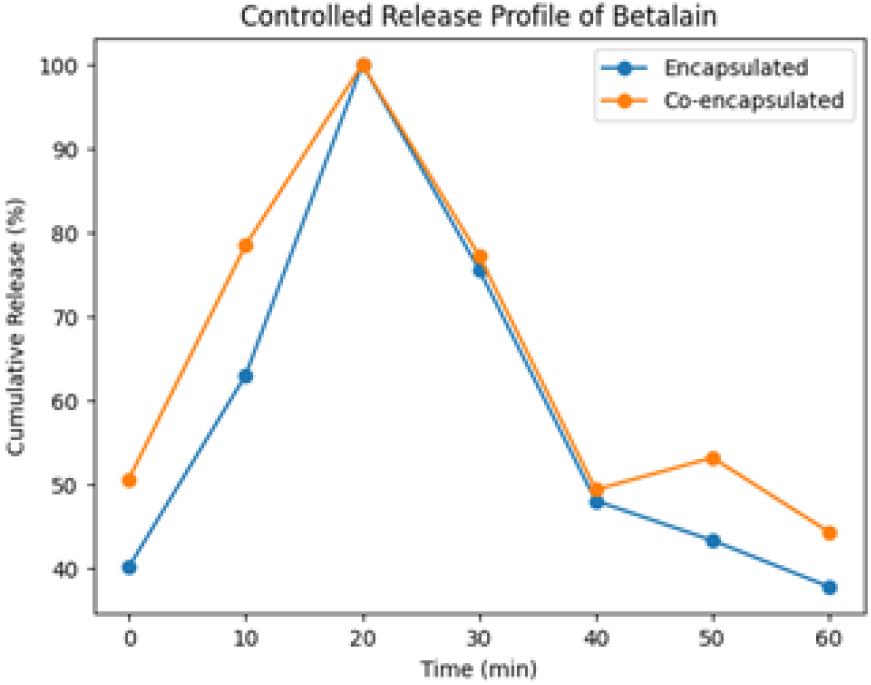
Controlled Release Profile

## V. Conclusion

In this work, betalain pigment was extracted from beetroot (Beta vulgaris) and quantified using UV–Visible spectrophotometry. The intrinsic instability of free betalain was greatly improved with alginate-based encapsulation and coencapsulation with food-grade materials. Encapsulation in calcium alginate beads enhanced pigment stability by reducing direct exposure to environmental stress factors: light, oxygen, and pH variations. Co-encapsulation of betalain with ascorbic acid provided better protection than encapsulation alone in improving color retention and pH tolerance under acidic conditions. The antioxidant activity of ascorbic acid together with the physical barrier exerted by the alginate matrix accounted for reduced oxidative degradation of the pigment. Controlled release studies further demonstrated that coencapsulated beads exhibited a more sustained and diffusion-controlled release profile than encapsulated beads, highlighting their potential for prolonged pigment delivery. Overall, the results have confirmed that alginate-based co-encapsulation is a very effective, sustainable, and food-compatible method for beetroot betalain stabilization [6], [18]. This approach will further widen the applicability of betalain, which is already a natural food colorant, to functionals foods and various controlled release delivery systems with clean label claims.

## VI. Applications and Future Scope

The improved stability properties and controlled release features of the encapsulated or co-encapsulated betalain pigments of the alginate-based encapsulation technology indicate its great potential for application in the food industry. The optimized stability properties ensure its appropriate usage for application as the natural food pigment for the coloring of liquid foods, candies, desserts, and other acidic food materials, providing a healthy alternative to synthetic food pigments [19], [20]. Aside from the color aspect, the antioxidant properties of betalain, together with the stability, allow the compound to be used in functional foods and supplements whose function is the delivery of health benefits [9], [11]. The controlled release properties of the co-encapsulated system are also advantageous in the formulation where the release of the pigment is needed. Betalain, which has sensitivity to pH, also has potential uses in smart food packaging, which could act as a natural pH indicator when betalain is encapsulated to detect food freshness and spoilage. These applications are in line with the need for smart and biodegradable food packaging materials [18]. Industrially, future research on the encapsulation of food materials should be concentrated on scale-up research to determine the viability of industrial production using extrusion or spray drying [15], [24]. Also, shelf-life research will be necessary to determine the viability of the encapsulated products. Additionally, research on other biopolymers, chitosan, gum arabic, and maltodextrin, may provide new means of further enhancing the shelf life and controlled release of food pigments. Advanced modeling of release rates may also provide new means of understanding the mechanisms of food pigment release and optimizing encapsulation systems.

